# Drosophila anion exchanger 2 is required for proper ovary development and oogenesis

**DOI:** 10.1101/459826

**Authors:** Marimar Benitez, Sumitra Tatapudy, Diane L. Barber, Todd Nystul

## Abstract

Understanding how cell fate decisions are regulated is a central question in stem cell biology. Recent studies have demonstrated that intracellular pH (pHi) dynamics contribute to this process. Indeed, the pHi of cells within a tissue is not simply a consequence of chemical reactions in the cytoplasm and other cellular activity, but is actively maintained at a specific setpoint in each cell type. We found previously that the pHi of cells in the follicle stem cell (FSC) lineage in the *Drosophila* ovary increases progressively during differentiation from an average of 6.8 in the FSCs, to 7.0 in newly produced daughter cells, to 7.3 in more differentiated cells. Two major regulators of pHi in this lineage are *Drosophila* sodium-proton exchanger 2 (*dNhe2*) and a previously uncharacterized gene, *CG8177*, that is homologous to mammalian anion exchanger 2 (AE2). Based on this homology, we named the gene *ae2*. Here, we generated null alleles of *ae2* and found that homozygous mutant flies are viable but have severe defects in ovary development and adult oogenesis. Specifically, we find that *ae2* null flies have smaller ovaries, reduced fertility, and impaired follicle formation. In addition, we find that the follicle formation defect can be suppressed by a decrease in *dNhe2* copy number and enhanced by the overexpression of *dNhe2*, suggesting that this phenotype is due to the dysregulation of pHi. These findings support the emerging idea that pHi dynamics regulate cell fate decisions and our studies provide new genetic tools to investigate the mechanisms by which this occurs.

## Introduction

The cell fate decisions that guide adult stem cell self-renewal and differentiation are controlled by a complex interplay of extracellular and intracellular signals. Many studies have focused on the role of cell signaling pathways in directing these cell fate decisions, but it is now becoming clear that other aspects of cell biology, such as changes in metabolism and cytoskeletal architecture, also contribute to this process. Consistent with this, we and others found that changes in intracellular pH (pHi) are important for adult and embryonic stem cell differentiation (Gao et al., 2014; Li et al., 2009; Ulmschneider et al., 2016). These findings reinforce the emerging view that changes in pHi are not only associated with pathological conditions such as cancer (White et al., 2017) and neurodegenerative disorders (Majdi et al., 2016), but also with normal cell behavior. For example, pHi has been observed to change across stages of the cell cycle (Putney and Barber, 2003), with cell migration and tissue remodeling (Jenkins et al., 2012; Stock and Schwab, 2009), and during cell differentiation and cell fate decisions (Tatapudy et al., 2017). Other studies have demonstrated that pHi dynamics function as a signaling mechanism by regulating “pH sensors”, defined as proteins with activities or binding affinities dependent on pH changes within cellular ranges (Schönichen et al., 2013). For example, increased pHi is necessary for the complex process of directed cell migration in part by regulating the pH sensors cofiin (Frantz et al., 2008), talin (Srivastava et al., 2008), and the focal adhesion kinase FAK (Choi et al., 2013). For pHi-dependent cell fate decisions, pH sensing by the histone H3 methylating enzyme Death Inducer Obliterator 3 (DIDO) (Tencer et al., 2017), metabolic enzymes (Tatapudy et al., 2017), and regulators of cell polarity (Frantz et al., 2007) are likely important. Hence, an area of current interest is identifying pHi-dependent cell behaviors and pHi-dependent processes regulating these behaviors. Moreover, because pHi dynamics are predominantly regulated by changes in the activity or expression of plasma membrane ion transport proteins (Casey et al., 2010), an important experimental direction is resolving how these ion transport proteins control cell behaviors.

To understand how ion transport proteins contribute to cell fate decisions in an adult epithelial stem cell lineage, we have been using the follicle stem cell (FSC) lineage in the *Drosophila* ovary as a model. Drosophila ovaries are composed of long strands of developing follicles, called ovarioles, and the FSCs reside in a structure at the anterior tip of each ovariole called the germarium (Margolis and Spradling, 1995). The FSCs divide with asymmetric outcomes during adulthood to self-renew and produce progeny, called prefollicle cells (pFCs). As new germ cell cysts are produced, they move past the FSCs and become surrounded by the pFCs, which then move along with the germ cell cysts through the germarium and differentiate into one of three major cell types, main body follicle cells, polar cells, or stalk cells. At the posterior end of the germarium, the cyst buds off, forming a round follicle with germ cells in the middle, a single layer of main body follicle cells around the outside, polar cells at the anterior and posterior poles, and stalk cells connecting one follicle to the next. The follicles then continue to grow and change shape as they progress through oogenesis. For reference, the germarium has been divided into four regions, 1, 2a, 2b and 3 (Fig. 1A), and follicles are classified into 14 stages, starting from the rounded cyst in Region 3 of the germarium as stage 1 and subsequent stages defined by size, shape and tissue morphology.

**Figure 1:**
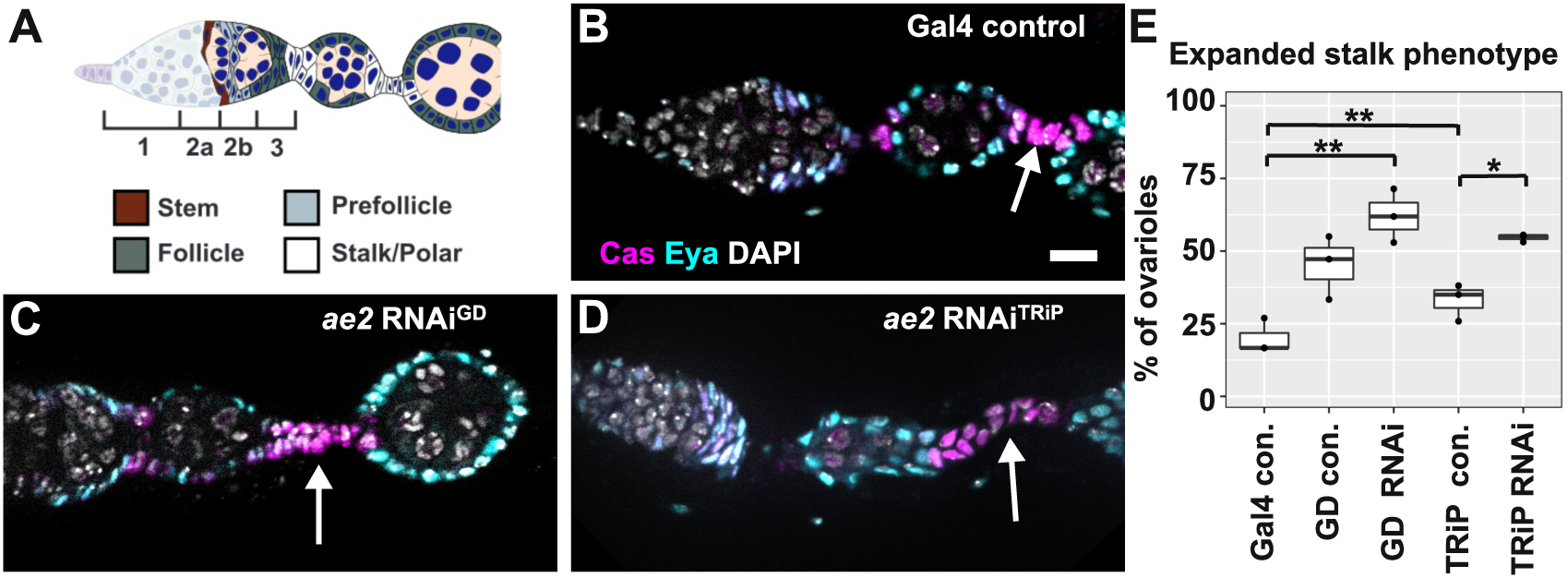
Identification of *ae2* in an RNAI screen. **(A)** Diagram of the germarium showing regions 1, 2a, 2b, and 3 of the germarium. FSCs (brown) are located in the middle of the germarium, at the region 2a/2b border. Newly produced daughter cells become pFCs (light gray) and then differentiate into main body FCs (dark gray), polar cells (white), or stalk cells (white). **(B-D)** A wildtype ovariole (B) or ovarioles with ae2 knockdown in follicle cells using *ae2^GD2397^* (C) or *ae2^HMJ30143^* **(D)** driven with *FC-Gal4* stained for Cas (magenta) to identify stalk cells, Eya (cyan) to identify main body follicle cells, and DAPI (white) to identify nuclel. **(E)** Graph showing the frequency of ovarioles with enlarged stalks in each of the indicated genotypes. “Gal4 con.” is *109-30-Gal4/CyO*; *tub-Gal80^ts^*, “GD con.” is *UAS-CG8177^GD2397^/CyO*, “GD RNAi” is *UAS-CG8177^GD2397^* combined with *109-30-Gal4*, “TRIP con.” is *UAS-CG8177^HMJ30143^/CyO* and “TRIP RNAI” is *UAS-CG8177^HMJ30143^* combined with *109-30-Gal4.* Scale bar represents 10 μm.* indicates p < 0.05 and ** indicates p < 0.01.

We previously reported that pHi increases during differentiation in the FSC lineage from an average of 6.8 in the FSCs, to 7.0 in pFCs, to 7.3 in main body follicle cells (Ulmschneider et al., 2016). We also found that the *Drosophila* sodium proton exchanger, *dNhe2*, is required for this increase in pHi, and that loss of *dNhe2* impaired pFC differentiation, particularly toward the stalk cell fate. Conversely, overexpression of *dNhe2* significantly raised pHi and caused an excess of cells in the stalk region. To further investigate a role for pHi in follicle cell differentiation, we performed an RNAi screen of 19 solute carrier (SLC) genes that are predicted to regulate pHi (Table S1), and assayed for follicle cell differentiation phenotypes that resembled loss or overexpression of *dNhe2*. We found that RNAi knockdown of *CG8177*, a previously uncharacterized gene, caused an increase in the pHi of follicle cells and an excess of cells in the stalk region, similar to the phenotype caused by *dNhe2*

Here, we present the full results of this screen and provide further characterization of the ovarian phenotypes caused by loss of *CG8177*, which we identify as encoding a Cl^−^/HCO_3_^−^ exchanger. We used CRISPR to generate two null alleles and observed multiple phenotypes in homozygous null flies. Specifically, we find that homozygous null flies are viable but have reduced fertility, fewer ovarioles, and several oogenesis phenotypes, including an excess of stalk cells, follicle encapsulation defects, and fewer follicles downstream from the germarium. Lastly, we found that these phenotypes in *CG8177* mutants are enhanced by *dNhe2* overexpression and suppressed by a reduction of *dNhe2*, suggesting that this phenotype is caused by dysregulation of pHi. Taken together, these findings provide support for the idea that pHi dynamics contribute to the control of cell fate decisions.

## Results

### *CG8177* is the homolog of mammalian anion exchanger 2

To identify genes that regulate pHi in the *Drosophila* ovary, we performed an RNAi-based screen through 19 genes that are expressed in the ovary according to the modENCODE tissue expression data available on FlyBase (Brown et al., 2014), and predicted to regulate pHi based on sequence homology (Table S1). We first combined RNAi lines for each gene with the follicle cell driver, *109-30-Gal4* (Hartman et al., 2010), which we refer to here as *FC-Gal4*, and *tub-Gal80^ts^* which allows for temporal control of expression. Then, we raised the progeny at 18°C to inhibit Gal4 activity, shifted to adults to 25°C for 7 days to allow for expression of the RNAi and knockdown the target gene, and screened for morphological phenotypes within the expression domain of *FC-Gal4*, which includes the follicle epithelium of the germarium and the first 2-3 budded follicles. We identified *CG8177* as the top candidate because RNAi knockdown caused one of the most reproducible and penetrant phenotypes. *CG8177* belongs to the SLC4A family of anion exchange proteins and has high homology with both the cytoplasmic and plasma membrane domains of members of the anion exhanger (AE) family (AE1-3) of mammalian Cl^−^/HCl3^−^ exchangers. A previous report referred to *CG8177* as *DAE*, for *Drosophila* anion exchanger (Dubreuil et al., 2010), though the gene is currently unnamed on FlyBase and more recent information available on FlyBase (Gramates et al., 2017) indicates that the closest reciprocal homolog within this family in both the mouse and human genomes is AE2, with 38% overall identity between the *Drosophila melanogaster* and human proteins, and 48% identity specifically within the anion exchange domain (Fig. S1). Therefore, we named the gene *anion exchanger 2* (*ae2*).

### Knockdown of *ae2* impairs stalk formation

*ae2* was identified in our previous screen because RNAi knockdown caused a significant increase in ovarioles with enlarged stalks. To confirm this RNAi phenotype, we tested two additional RNAi lines and again found a significant increase in ovarioles with enlarged stalks between budded follicles, from 20 ± 6% in the *FC-Gal4* control to 62 ± 9% and 55 ± 1% in flies with *FC-Gal4* driving expression of *ae2^GD2397^* or *ae2^HMJ30143^*, respectively (Fig 1B-E). This phenotype was rare in wildtype (Oregon-R) flies, but occurred with moderate penetrance in the RNAi lines alone (without the *FC-Gal4* driver, Fig. 1E), and the penetrance did not diminish after multiple generations of outcrossing to Oregon-R. These data suggest that the phenotypes in the RNAi lines are due to leaky expression of the RNAi rather than background mutations in the stocks. Several other RNAi lines in the screen also exhibited a stalk phenotype with moderate penetrance in the absence of the driver, including CG32081 and CG1139, and the penetrance increased when the RNAi was driven with *FC-Gal4*. However, unlike the *ae2* lines, the difference in penetrance with and without the driver was not significant, so they were not prioritized in this study.

### Loss of *ae2* impairs ovarian development and female fertility

To circumvent technical issues with the RNAi lines and to test *ae2* function more broadly, we used CRISPR (Bassett and Liu, 2014; Gratz et al., 2014) to introduce small deletions in two target sites within the first coding exon that is common to all the annotated isoforms of *ae2* (Fig. S2). We obtained one line with a 7 base pair deletion in Target 1 (*ae2^1^*), and another line with a 13 base pair deletion in Target 2 (*ae2^2^*). Both mutations cause a frameshift near the 5’ end of the coding region that results in an early stop codon, and thus are expected to be null alleles. Homozygous *ae2* null flies survived to adulthood, and one of the first phenotypes we noticed was that the ovaries were substantially smaller in homozygotes compared with heterozygotes or the *Cas9* parental stock that was used to generate the CRISPR alleles (Fig. 2A-C). To further investigate this phenotype, we quantified the number of ovarioles per ovary pair in young flies (less than 3 days post eclosion) and found that *ae2^1^* homozygotes had significantly fewer ovarioles than their heterozygous siblings or the *Cas9* parental stock (Fig. 2D, 13 ± 4 versus 20 ± 6 and 22 ± 4 respectively). Ovarioles are already fully formed when adults eclose from the pupal case, so this finding suggests that *ae2* functions during ovarian development in the process of ovariole specification. Consistent with the reduced ovariole number in *ae2^1^* homozygotes, we found a similar reduction in the number of eggs deposited per day by *ae2^1^* homozygotes compared to either a wildtype stock containing the same balancers or the parental *Cas9* control (Fig. 2E). Taken together, these data indicate that loss of *ae2* impairs ovarian development and female fertility.

**Figure 2:**
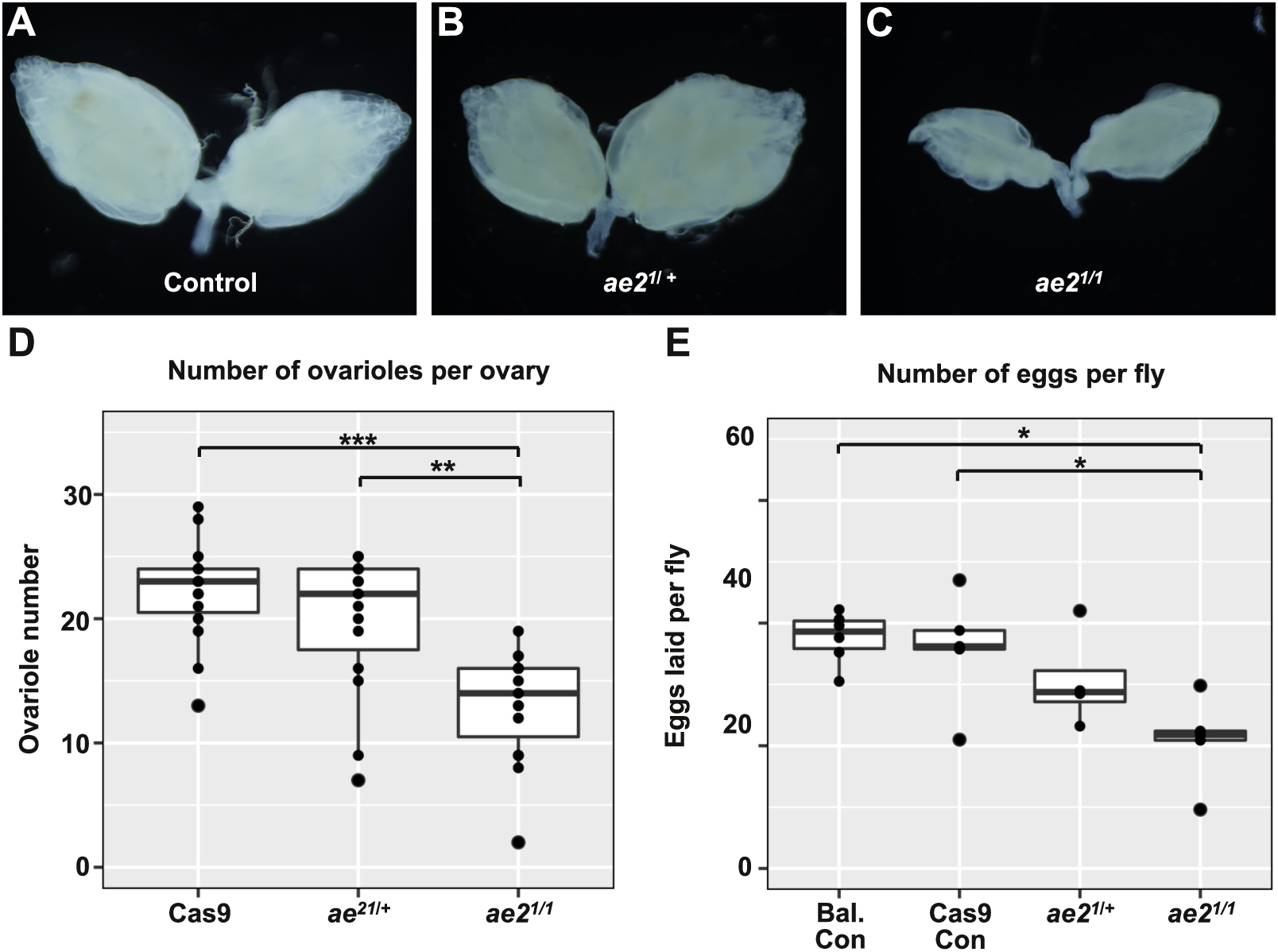
*ae2* mutants have reduced ovary size and fertility. **(A-C)** Ovaries from wildtype (A), *ae2*^1/^ + heterozygotes (B), or *ae2^1^/ae2^1^* homozygotes (C). Ovaries from homozygous flies are much smaller than in wildtype or heterozygous flies. **(D-E)**Graphs showing the number of ovarioles per ovary pair (D) and the number of eggs laid per day per female (E) for the indicated genotypes. Bal con. is *sp/CyO*; *TM2/TM6* and Cas9 con. is *nos-Cas9/CyO*. * indicates p < 0.05, ** indicates p < 0.01, and *** indicates p < 0.001.

### Loss of *ae2* impairs follicle development

We next examined individual ovarioles in *ae2* homozygous nulls and found several follicle formation defects, including an enlarged stalk that resembled the phenotype caused by expression of *ae2* RNAi in follicle cells, a “fusion” phenotype, in which multiple germ cell cysts were packaged into the same follicle layer, and small cysts with apoptotic germ cells (Fig. 3A-H). Ovarioles with these phenotypes invariably had fewer follicles between the germarium and late stage follicles (≥ stage 10), likely because these defects prevented the formation, development or survival of newly budded follicles (Fig. 3E-H). In some cases, no defective follicles were present but ovarioles still had fewer follicles, perhaps because the follicle formation defects slowed the rate of budding or the defective follicles had already degraded. In the *Cas9* parental stock and *ae2^1^*/+ heterozygotes, these follicle formation defects were rare. As expected, most ovarioles (94 ± 3% in the *Cas9* stock and 95 ± 2% in ae21/+ heterozygotes) had a normal complement of follicles (> 4 follicles) downstream from the germarium that were distributed among the early, mid, and late stages of oogenesis (Fig. 3A-B and J). In contrast, 55% ± 8% of *ae2^1^* and 49 ± 16% of *ae2*^2^ homozygous ovarioles contained fewer than four follicles between the germarium and the first late stage follicle (Fig. 3E-G and J). We observed a similar penetrance of this phenotype in *ae2^1^*/*ae2^2^* transheterozygotes (Fig. 3H and J), and found that the penetrance in an *ae2^1^* homozygous background was significantly reduced by the addition of a wildtype *ae2* transgene (Fig. 3I-J), indicating that the phenotype is due to the loss of *ae2* function in these mutants. These findings indicate that loss of *ae2* severely disrupts follicle development.

**Figure 3:**
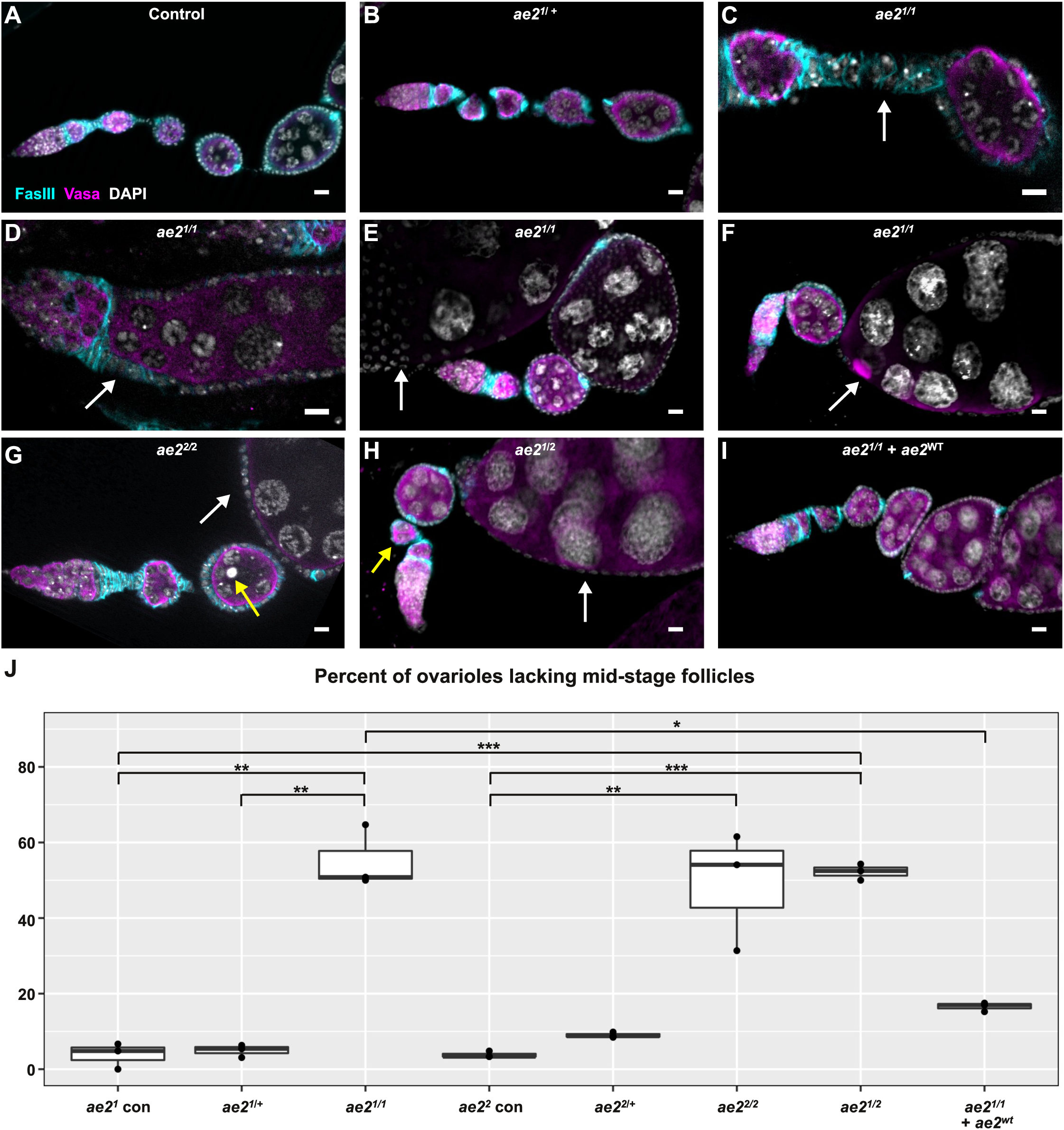
*ae2* is required for proper oogenesis. **(A-I)** Ovarioles from *Cas9* control flies (A), *ae2^1^*/ + heretozygotes (B), *ae2^1^* homozygotes (C-F), *ae2^2^* homozygotes (G), *ae2^1^/ae2^2^* transheterozygotes (H) or *ae2^1^* homozygotes rescued with a *ae2^wt^* transgene (I) stained for Faslll (cyan) to identify follicle cells vasa (magenta) to identify germ cells, and DAPI (white) to identify nuclei. *ae2* null mutants have several phenotypes, including enlarged stalks (C), fused egg chambers with multiple germ cell cysts packeged into a single follicle epithelium (D, white arrow), apoptotic germ cells (G, yellow arrow), abnormally small follicles (H, yellow arrow), and a reduced number of mid-stage follicles between the germarium and late stage follicles (E-H, white arrows indicate late stage follicles). **(J)** Graph showing the frequency of ovarioles with reduced numbers of mid-stage follicles for the indicated genotypes. *ae2^1^* con. and *ae2^2^* con. are *Cas9* controls that were cultured and processed in parallel with the respective mutant genotypes. Scale bar represents 10 μm. * indicates p < 0.05, ** indicates p < 0.01, and *** indicates p < 0.001. Not shown on the graph: the frequency of ovarioles with reduced numbers of mid-stage follicles in *ae2^1^* homozygotes (from Figure 3) is significantly higher than in *dNhe2^null^/+; ae2^1^* (p < 0.01).

### *ae2* genetically interacts with *dNhe2* in oogenesis

SLC4A family members function as “acid loaders” by catalyzing an efflux of intracellular HCO_3_^−^ to decrease pHi (Alper, 2006; Casey et al., 2010), so loss of *ae2* would be predicted to cause an increase in pHi. Indeed, we previously found that pHi significantly increases with RNAi knockdown of *ae2* in follicle cells (Ulmschneider et al., 2016). Conversely, *dNhe2* functions to increase pHi by extruding protons, and RNAi knockdown of *dNhe2* in follicle cells significantly decreases pHi (Ulmschneider et al., 2016). Therefore, if the phenotypes in *ae2* null homozygotes are due to dysregulation of pHi, we reasoned that they should be enhanced by overexpression of *dNhe2* and suppressed by a decrease in *dNhe2* function. To test this prediction, we first quantified the follicle cell development phenotype in *ae2^1^* heterozygotes or homozygotes in combination with *dNhe2* overexpression in follicle cells. Indeed, although the frequency of the follicle development phenotype was low in *ae2^1^* heterozygotes (5 ± 2%, Fig. 3B and J) and in flies with *FC-Gal4* driving *dNhe2* overexpression alone (3 ± 4%, Fig 4A and G), we found that the frequency of the phenotype was significantly increased in *ae2^1^* heterozygotes combined with *dNhe2* driven by *FC-Gal4* (45 ± 13%, Fig. 4B and G). Notably, overexpression of *dNhe2* did not further enhance the phenotype in *ae2^1^* homozygotes. To test for a genetic interaction in the opposite direction, we combined *ae2^1^* heterozygotes or homozygotes with heterozygosity for *dNhe2^null^*. The frequency of the follicle development phenotype was low in the *dNhe2^null^*/ +;*ae2^1^*/ + double heterozygotes (12 ± 7%, Fig. 4E and G), as expected, and not significantly different than the *Cas9* control (8 ± 4%, Fig. 4D). Interestingly, removal of one copy of *dNhe2* significantly reduced the frequency of the follicle development phenotype in *ae2^1^* homozygotes to a value that was not significantly different from the *Cas9* control (20 ± 8% in *dNhe2^null^*/+; *ae2^1^* vs 55 ± 8% in *ae2^1^* and 8 ± 4% in *Cas9* controls, Fig. 4D, F, and G). These genetic interactions between *dNhe2* and *ae2* suggest that the follicle development phenotype in *ae2* null mutants is caused by dysregulation of pHi.

**Figure 4:**
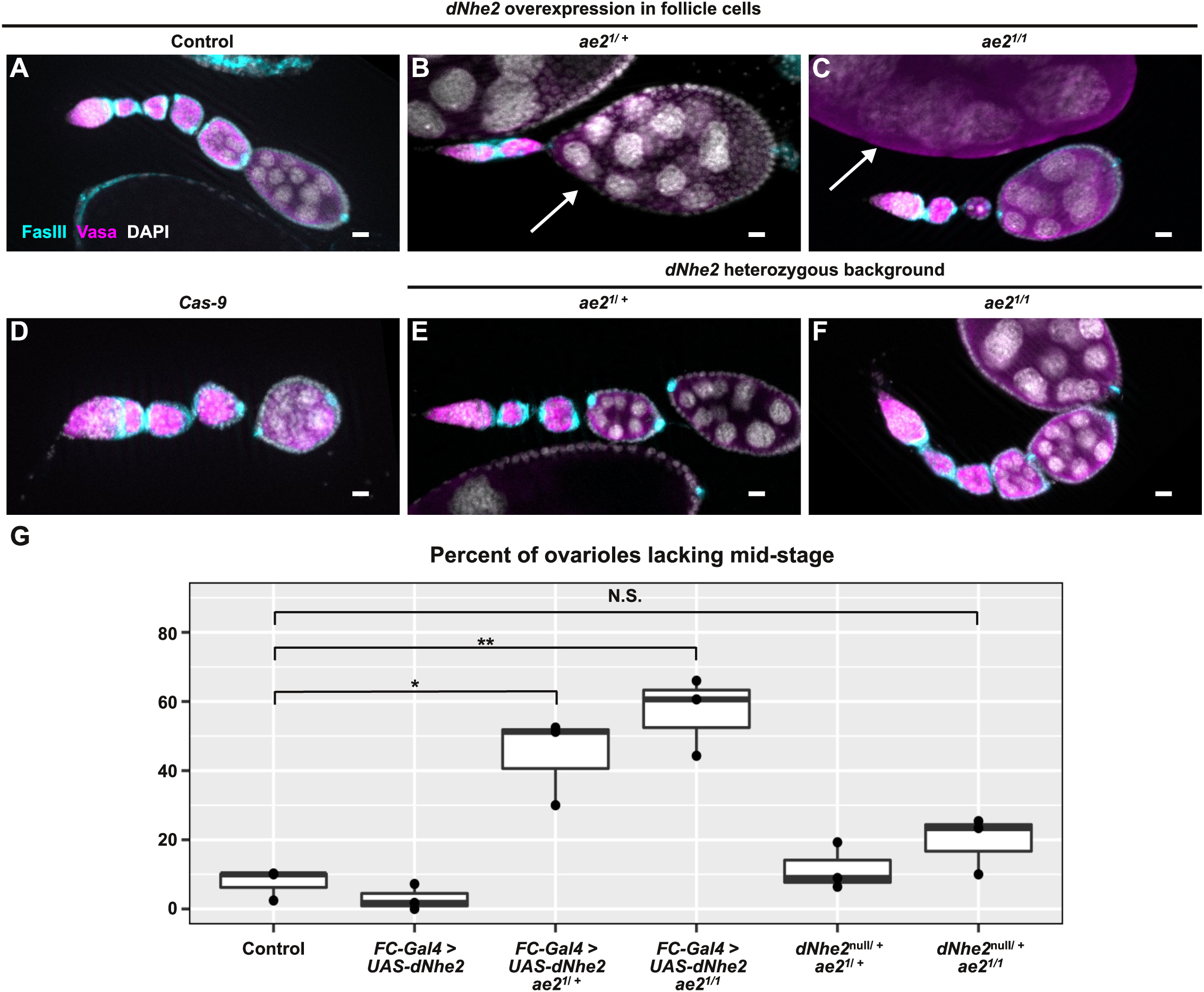
*ae2* genetically interacts with *dNhe2*. **(A-C)** Ovarioles from flies with overexpression of *dNhe2* in follicle cells alone as a control (A) or in combination with *ae2^1^*/ + (B) or *ae2^1^* (C). Overexpression of *dNhe2* does not affect the presence of mid-stage follicles in a wildtype background, but causes a reduction in mid-stage follicles in combination with *ae2^1^*/+ heterozygosity. **(D-F)**Ovarioles from a *Cas9* control (D) or *dNhe2/+* flies combined with *ae2^1^*/+ (E), or *ae2^1^/ae2^1^* (F). Heterozygosity for *dNhe2* rescues the mid-stage follicle phenotype in *ae2^1^* homozygotes. **(G)** Graph showing the frequency of ovarioles with reduced numbers of mid-stage follicles for the indicated genotypes. Control is *nos-Cas9/CyO*. Ovarioles are stained for Faslll (cyan) to identify follicle cells, vasa (magenta) to identify germ cells, and DAPI (white) to identify nuclei. Scale bar represents 10 μm. * indicates p < 0.05, ** indicates p < 0.01, and N.S. indicates Not Significant.

## Discussion

In this study, we characterized the phenotypes in *Drosophila* ovary development and follicle formation caused by loss of *ae2* (previously referred to as *DAE* or *CG8177*). We first identified the role for *ae2* in ovaries by an RNAi screen through SLC genes for phenotypes in follicle cell development, and found that RNAi knockdown of *ae2* caused enlarged stalks. Here, we generated null alleles of *ae2* and found that homozygous null ovarioles have not only the enlarged stalk phenotype but also several other more severe phenotypes, including a follicle fusion phenotype in which multiple germ cell cysts are packaged together into the same follicle and germ cell apoptosis. These phenotypes are very likely to be incompatible with healthy follicle development and, indeed, we found that the most common feature of mutant ovarioles was a significant reduction in the number of follicles per ovariole. We found previously that *dNhe2* and *ae2* regulate pHi in the follicle epithelium (Ulmschneider et al., 2016), but these proteins also transport other ions in the process (sodium in the case of *dNhe2* and chloride in the case of *ae2*), so the loss-of-function phenotypes could be due, in part, to an imbalance of these other ions. However, our finding that the follicle formation phenotypes in *ae2* mutants can be suppressed by a reduction in *dNhe2* copy number and enhanced by overexpression of *dNhe2* strongly suggest that these phenotypes are due to dysregulated pHi. This is consistent with our previous findings that pHi dynamics are important for cell fate specification in the early FSC lineage (Ulmschneider et al., 2016).

Additionally, our findings that ovary size and ovariole number are substantially reduced in newly eclosed *ae2* homozygous nulls suggest that *ae2* also functions during ovary development. Ovarioles are established during larval development by the differentiation of intermingled cells into either terminal filament cells or basal stalk cells. These cell fate specifications are controlled by hippo (Sarikaya and Extavour, 2015) and hedgehog signaling (Lai et al., 2017), and the process by which the cells coalesce into rudimentary ovarioles requires cell migration, intercalation (Godt and Laski, 1995; Li et al., 2003), and actomyosin contractility (Vlachos et al., 2015). Interestingly, several of these processes are influenced by pH dynamics in other contexts. Specifically, dysregulation of pHi affects the pattern of Hedgehog signaling in the adult *Drosophila* ovary (Ulmschneider et al., 2016), and actin filament assembly and cell migration in mammalian fibroblasts (Denker and Barber, 2002; Frantz et al., 2008, 2007). In addition, adherens junction remodeling is important for cell intercalation, and β-catenin, which is a core component of adherens junctions, is also pH sensitive (White et al., in press). Therefore, it will be interesting to investigate whether these potentially pH-sensitive steps are responsible for the ovarian development phenotypes we observed in *ae2* homozygous null flies.

Our findings in this study align well with studies of pHi during mammalian oogenesis. Similar to our finding that *dNhe2* and *ae2* are important regulators of pHi in the *Drosophila* ovary, anion exchangers and sodium proton exchangers are also known to be the primary means of regulating pHi in mammalian ovaries (FitzHarris and Baltz, 2009). Interestingly, mammalian germ cells in early oogenesis are unable to regulate pHi on their own and instead rely on the activity of anion exchangers and sodium proton exchangers in the surrounding granulosa cells (FitzHarris et al., 2007; Fitzharris and Baltz, 2006). These cells are thought to regulate germ cell pHi directly by transporting protons through gap junctions. Follicle cells in the *Drosophila* ovary are the functional equivalent of granulosa cells in the mammalian ovary, and as with granulosa cells, *Drosophila* follicle cells are connected to germ cells through gap junctions (Bohrmann and Zimmermann, 2008; Mukai et al., 2011; Tazuke et al., 2002). Thus, the loss of *ae2* in follicle cells may cause follicle development defects at least in part through a dysregulation of germ cell pHi. Although the mechanism by which dysregulation of pHi causes impaired follicle development is not known, a recent study of ovarian cancer cell growth suggests that one point of regulation may be mTOR (Zhang et al., 2017), which is a well-known regulator of cell growth and proliferation. This study found that mTOR pathway activity was associated with AE2 expression, and that the effects of mTOR inhibition on cell cycle progression could be mitigated by overexpression of AE2. In the FSC lineage, the *Drosophila* homolog of mTOR, *Tor*, regulates proliferation of FSCs but not the downstream differentiating progeny (LaFever et al., 2010). Thus, one role for pHi dynamics in the early FSC lineage may be to regulate *Tor* activity.

Collectively, our data provide additional support for the idea that pHi dynamics play an important role in the regulation of developmental and physiological processes, and introduce new mutant alleles that are useful for manipulating pHi *in vivo*. Our studies with these alleles highlight stages of ovarian development and oogenesis that are sensitive to dysregulated pHi dynamics and provide a foundation for future studies into the molecular mechanisms by which pHi contributes to the regulation of cell fate.

## Materials and Methods

### Fly Stocks

Stocks were maintained on standard molasses food at 25°C and adults were given fresh wet yeast daily. All progeny containing tub-Gal80^ts^ were kept at 18°C until eclosion and then shifted to 29oC for 7-10 days for high UAS expression. The following stocks were used.

1. y1, w*; P{GawB}109-30/CyO; (BL#7023)
2. w1118; P{GD2397}v39492/TM3 (VDRC#39492)
3. y[1] v[1]; P{y[+t7.7] v[+t1.8]=TRiP.HMJ30143}attP40 (BL#63577)
4. w; 10930/Cyo; tub-Gal80^ts^
5. *y^2^ cho2 v1*; attP40{nos-Cas9}/CyO (Rainbow Transgenic, CAS-0001)
6. *w*; *sp*/CyO; *ae2^1^*/TM2 (Nystul Lab)
7. *w*; *sp*/CyO; *ae2^2^*/TM2 (Nystul Lab)
8. ; CG8177^WT^ (Genetivision, stock #P9-C9)
9. OregonR (BL#5)
10. *w*;*109-30*, *UAS-DNhe2*/*CyO*; *tub-Gal80^ts^*/TM2 (Barber Lab)
11. *w; dNhe2^nul^* (Barber Lab)

### Immunostaining

Ovaries were dissected in 1x phosphate buffered saline (PBS), fixed in 1x PBS + 4% formaldehyde for 15 minutes, rinsed with 1x PBS + 0.2% Triton X-100 (PBST) and blocked for 1 hour with 1x PBST containing 0.5% BSA. Samples were incubated with primary antibodies diluted in blocking solution overnight at 4 deg C. Next, samples were rinsed with PBST and blocked for 1 hour before incubating with secondary antibodies for 2 hours at room temperature. Samples were rinsed twice with PBST and once with PBS before a final 30 minute wash with PBS. Samples were mounted on glass slides in Vectashield (Vector Labs) with DAPI. The following primary antibodies were used: guinea pig anti GFP [1:1000] (Synaptic Systems 132005), rabbit anti-Vasa [1:1000] (Santa Cruz sc-30210), mouse anti-armadillo [1:100] (DSHB N27A1), mouse anti-Eyes absent [1:50] (DSHB 10H6), rabbit anti-Castor [1:5000] (from Ward Odenwald) (Kambadur et al., 1998). The following secondary antibodies were purchased from Thermo Fisher Scientific and used at 1:1000: goat anti-guinea pig 488 (A-11073), goat anti rabbit 488 (A-11008), goat anti rabbit 555 (A-21428), goat anti mouse 488 (A-11029), goat anti-mouse 555 (A-21424). All fixed images were acquired using a Zeiss M2 Axioimager with Apotome unit. For multicolor fluorescence images, each channel was acquired separately. Post acquisition processing such as image rotation, cropping, and brightness or contrast adjustment were performed using ImageJ and Photoshop. Acquisition settings and any brightness/contrast adjustments were kept constant across conditions within an experiment.

### Egg laying assay

1-2 day old females of the indicated genotypes were collected and fed wet yeast for 2 consecutive days in the presence of wild type males. 5 females together with 3 males were then allowed to deposit eggs for 24 hours on molasses plates with yeast at 25 °C. Data is presented as number of eggs deposited per female.

### Statistics

Statistics were performed using R Studio. An R Notebook showing statistical procedures and outputs, including p-values and graphs, is supplied as a supplemental file.

## Acknowledgements

We thank Katja Rust for performing the egg laying assay, the Bloomington Stock Center and the Vienna Drosophila Resource Center for stocks, and FlyBase for information and analysis. This work was funded by National Institute of Health grant GM116384 to DLB and TGN.

---

**Figure S1:**
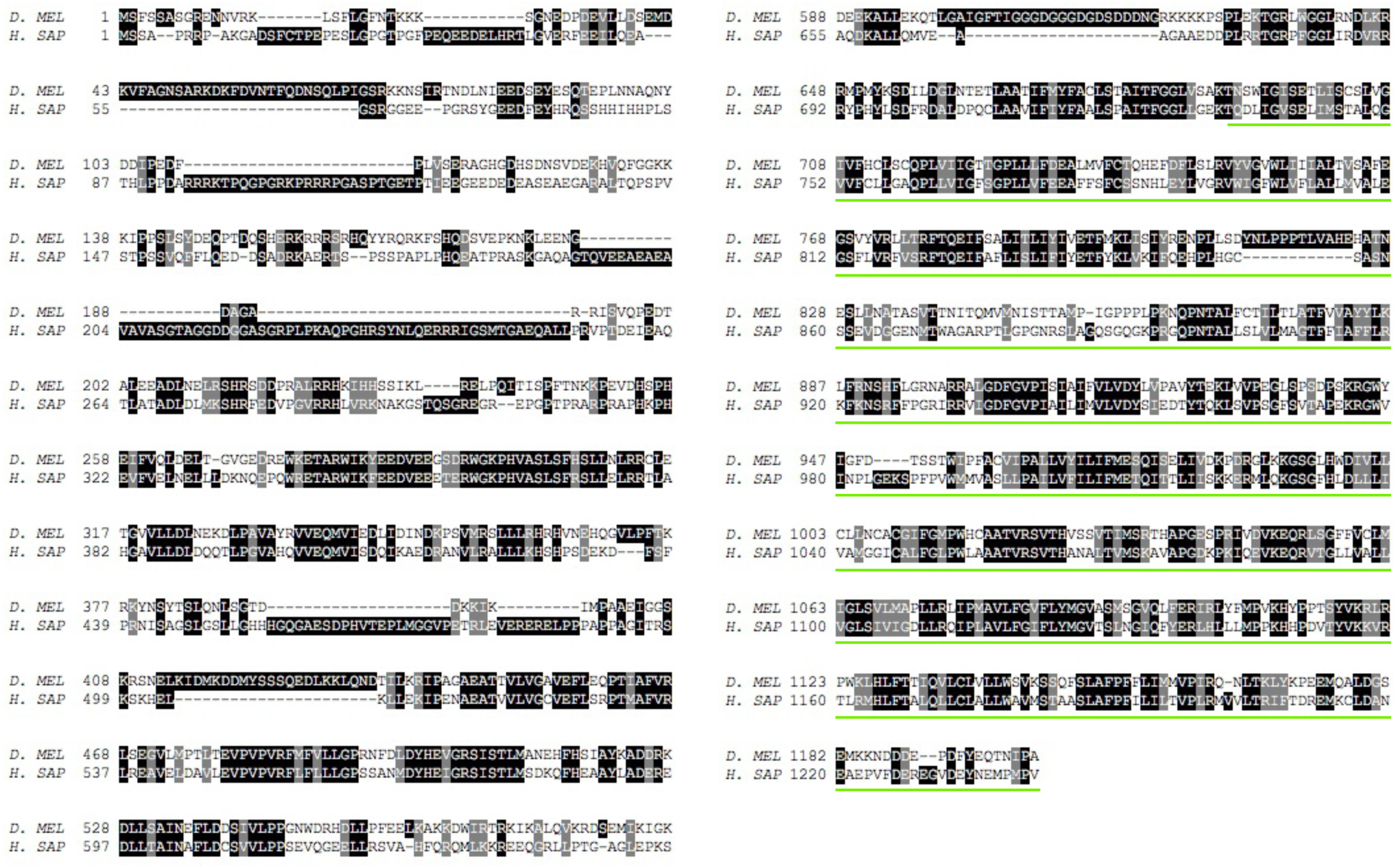
Alignment of the Drosophila and human AE2 genes. Protein alignment from Drosohila and human *ae2* genes. The anion exchange domain is Indicated by the green bars.

**Figure S2:**
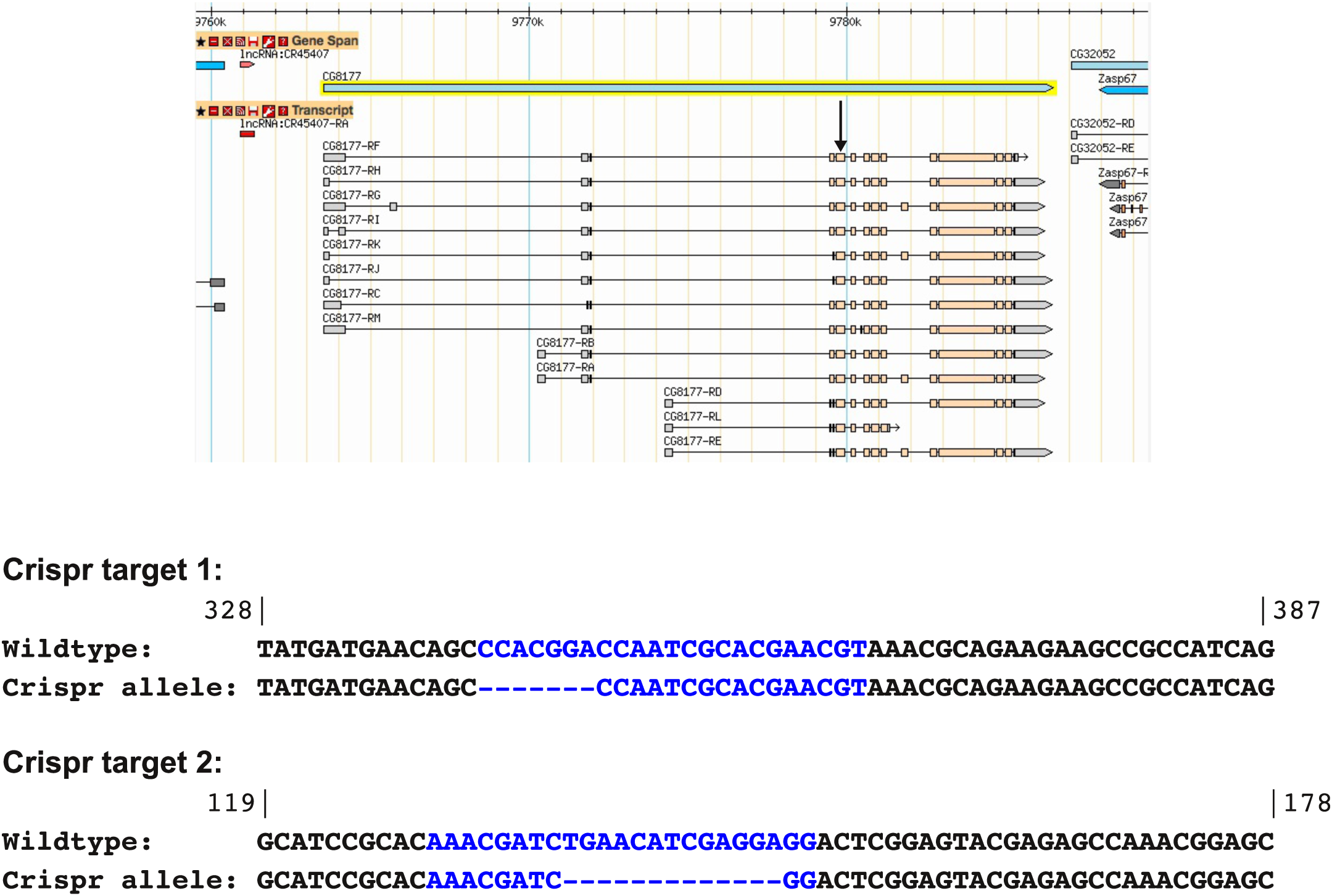
Generation of *ae2* CRISPR allels. (A) Screen shot from GBrowse on FlyBase showing the annotated mRNA isoforms of *ae2.* (B) Sequences within *ae2* that were targeted by CRISPR. Target sequences shown in blue and flanking sequences shown in black. The alleles recovered had deletions as indicated.

**Table S1:** RNAi screen of SLC genes that are expressed in the ovary and predicted to regulate pHi.

| Gene name | Genotype | Stock number | Phenotype<sup>1</sup> | Function | Predicted effect on pH<sub>i</sub> |
| --- | --- | --- | --- | --- | --- |
| CG8177 | *UAS-CG8177 RNAi[GD2397]* | VDRC#39492 | Yes | Cl/HCl<sub>3</sub> exchanger | Increases |
| CG8177 | *UAS-CG8177 RNAi[HMJ30143]* | BL#63577 | Yes | Cl/HCl<sub>3</sub> exchanger | Increases |
| CG8785 | *UAS-CG8785 RNAi[GD1961]* | VDRC#4650 | No | Amino acid transporter | Increases |
| CG8785 | *UAS-CG8785 RNAi[GD1961]* | VDRC#4651 | No | Amino acid transporter | Increases |
| CG32081 | *UAS-CG32081 RNAi[GD2410]* | VDRC#5209 | Yes | Amino acid transporter | Increases |
| CG32081 | *UAS-CG32081 RNAi[KK105324]* | VDRC#107023 | Yes | Amino acid transporter | Increases |
| Kar | *UAS-Kar RNAi[KK101781]* | VDRC#105429 | No | Monocarboxylate transporter | Decreases |
| Path | *UAS-Path RNAi[GD2391]* | VDRC#44536 | No | Amino acid transporter | Increases |
| Path | *UAS-Path RNAi[GD2391]* | VDRC#44537 | No | Amino acid transporter | Increases |
| Sln | *UAS-Sln RNAi[GD1940]* | VDRC#4607 | No | Monocarboxylate transporter | Decreases |
| Sln | *UAS-Sln RNAi[GD1940]* | VDRC#4609 | No | Monocarboxylate transporter | Decreases |
| Sln | *UAS-Sln RNAi[KK104306]* | VDRC#109464 | No | Monocarboxylate transporter | Decreases |
| CG1139 | *UAS-CG1139 RNAi[GD3830]* | VDRC#8907 | No | Amino acid transporter | Increases |
| CG1139 | *UAS-CG1139 RNAi[KK111566]* | VDRC#102363 | No | Amino acid transporter | Increases |
| Nhe3 | *UAS-Nhe3 RNAi[GLC01631]* | BL#50513 | No | Na<sup>+</sup>/H<sup>+</sup> exchanger | Decreases |
| CG16700 | *UAS-Nhe3 RNAi[KK105945]* | VDRC#110058 | Yes | Amino acid transporter | Increases |
| CG12943 | *UAS-CG12943 RNAi[KK112469]* | VDRC#107119 | Yes | Amino acid transporter | Increases |
| CG32079 | *UAS-CG32079 RNAi[KK107121]* | VDRC#104454 | Yes | Amino acid transporter | Increases |
| Calx | *UAS-Calx RNAi[JF02937]* | BL#28306 | No | Na/Ca-exchange protein | Decreases |
| CG11665 | *UAS-CG11665 RNAi[HMC03642]* | BL#52902 | No | monocarboxylate transporter | Decreases |
| CG13384 | *UAS-CG13384 RNAi[HMS02268]* | BL#41703 | Yes | amino acid transporter | Increases |
| CG4991 | *UAS-CG4991 RNAi[GLC01599]* | BL#44450 | No | amino acid transporter | Increases |
| CG7888 | *UAS-CG7888 RNAi[GD2411]* | VDRC#37263 | No | amino acid/proton symporter | Increases |
| CG7888 | *UAS-CG7888 RNAi[GD2411]* | VDRC#37264 | No | amino acid/proton symporter | Increases |
| Ndae1 | *UAS-Ndae1 RNAi[GD967]* | VDRC#3664 | No | Na<sup>+</sup> anion exchanger | Decreases |
| Ndae1 | *UAS-Ndae1 RNAi[KK100362]* | VDRC#107948 | No | Na<sup>+</sup> anion exchanger | Decreases |
| Nhe1 | *UAS-Nhe1 RNAi[HM05077]* | BL#28589 | No | Na<sup>+</sup>/H<sup>+</sup> exchange | Decreases |
| CG43693 | *UAS-CG43693 RNAi[GL01304]* | BL#41873 | No | Amino acid transporter | Increases |
| CG43693 | *UAS-CG43693 RNAi[GLV21061]* | BL#35696 | No | Amino acid transporter | Increases |

1. Indicates whether a follicle cell phenotype was observed in the follicle epithelium when RNAi line was combined with FC-Gal4. However, CG8177 RNAi lines were the only ones in which the penetrance of the phenotype was significantly higher than in the parental RNAi line alone.

